# Effect of inbreeding on type 2 diabetes-related metabolites in a Dutch genetic isolate

**DOI:** 10.1101/618801

**Authors:** Ayşe Demirkan, Jun Liu, Najaf Amin, Ko Willems van Dijk, Cornelia M. van Duijn

**Author notes:** These authors contributed equally (Ayşe Demirkan, Jun Liu). **Materials and Correspondence:** Correspondence and requests for materials should be addressed to Ayşe Demirkan or to Jun Liu or to or to Cornelia M. van Duijn.

## Abstract

Autozygosity, meaning inheritance of an ancestral allele in the homozygous state is known to lead bi-allelic mutations that manifest their effects through the autosomal recessive inheritance pattern. Autosomal recessive mutations are known to be the underlying cause of several Mendelian metabolic diseases, especially among the offspring of related individuals. In line with this, inbreeding coefficient of an individual as a measure of cryptic autozygosity among the general population is known to lead adverse metabolic outcomes including type 2 diabetes (T2DM), a multifactorial metabolic disease for which the recessive genetic causes remain unknown. In order to unravel such effects for multiple metabolic facades of the disease, we investigated the relationship between the excess of homozygosity and the metabolic signature of T2DM. We included a set of heritable 143 circulating markers associated with fasting glucose in a Dutch genetic isolate Erasmus Rucphen Family (ERF) of up to 2,580 individuals. We calculated individual whole genome-based, exome-based and pedigree-based inbreeding coefficients and tested their influence on the T2DM-related metabolites as well as T2DM risk factors. We also performed model supervised genome-wide association analysis (GWAS) for the metabolites which significantly correlate with inbreeding values. Inbreeding value of the population significantly and positively correlated with associated with risk factors of T2DM: body-mass index (BMI), glucose, insulin resistance, fasting insulin and waist-hip ratio. We found that inbreeding influenced 32.9% of the T2DM-related metabolites, clustering among chemical groups of lipoproteins, amino-acids and phosphatidylcholines, whereas 80 % of these significant associations were independent of the BMI. The most remarkable effect of inbreeding is observed for S-HDL-ApoA1, for which we show evidence of the novel *DISP1* genetic region discovered by model supervised GWAS, in the ERF population. In conclusion, we show that inbreeding effects human metabolism and genetic models other than the globally used additive model is worth considering for study of metabolic phenotypes.

## Introduction

Autozygosity, meaning inheritance of an ancestral allele in the homozygous state is known to lead bi-allelic mutations that manifest their effects through the autosomal recessive inheritance pattern. Offspring of related individuals are at an increased risk of inheriting two copies of recessive deleterious alleles, which would expose the offspring to the full (normally compensated) deleterious effects of those alleles, hence decreasing the fitness of the offspring. Consanguineous marriages between close relatives as a result of assertive mating is known to cause severe congenital metabolic consequences in the off-spring^1^. In addition to that moderate inbreeding due to isolation in populations has been shown to cause unfavorable outcomes among with cardio-metabolic and neuropsychiatric parameters^2,3^. Inbreeding was shown to associate with an increase in fasting glucose, blood pressure, body mass index (BMI), waist-hip ratio (WHR) and decrease in high-density lipoprotein cholesterol (HDL-C), intelligence quotient (IQ) and height^4,5^. We have previously shown that some metabolites can well be regulated by genetic variants following the recessive genetic model^6^. Although most of the inborn errors of metabolism are inherited via the recessive genetic model requiring both copies of the metabolic enzymes to be impaired, metabolic outcomes under the control of cryptic relatedness have not been investigated thoroughly at the population level.

Recent technological advances in metabolomics allow us to capture the current state of biochemical events in an organism. The metabolome level is the result of our genome and environmental exposures. Application of metabolomics to metabolic disorders, especially type 2 diabetes (T2DM), is particularly promising since deregulations of biochemical processes are involved in the pathophysiology of disease^7,8^. In line with this, several circulating molecules have been found associated with T2DM: such as phospholipids, branch-chain amino-acids and lipoprotein subclasses^9–11^. In order to find target endophenotypes for researching the recessive genetic effects, we studied the influence of inbreeding over a selected list of metabolites that are known to be related to changes in glucose. **Figure 1** shows the outline of the step-wise analysis design.

**Figure 1.**
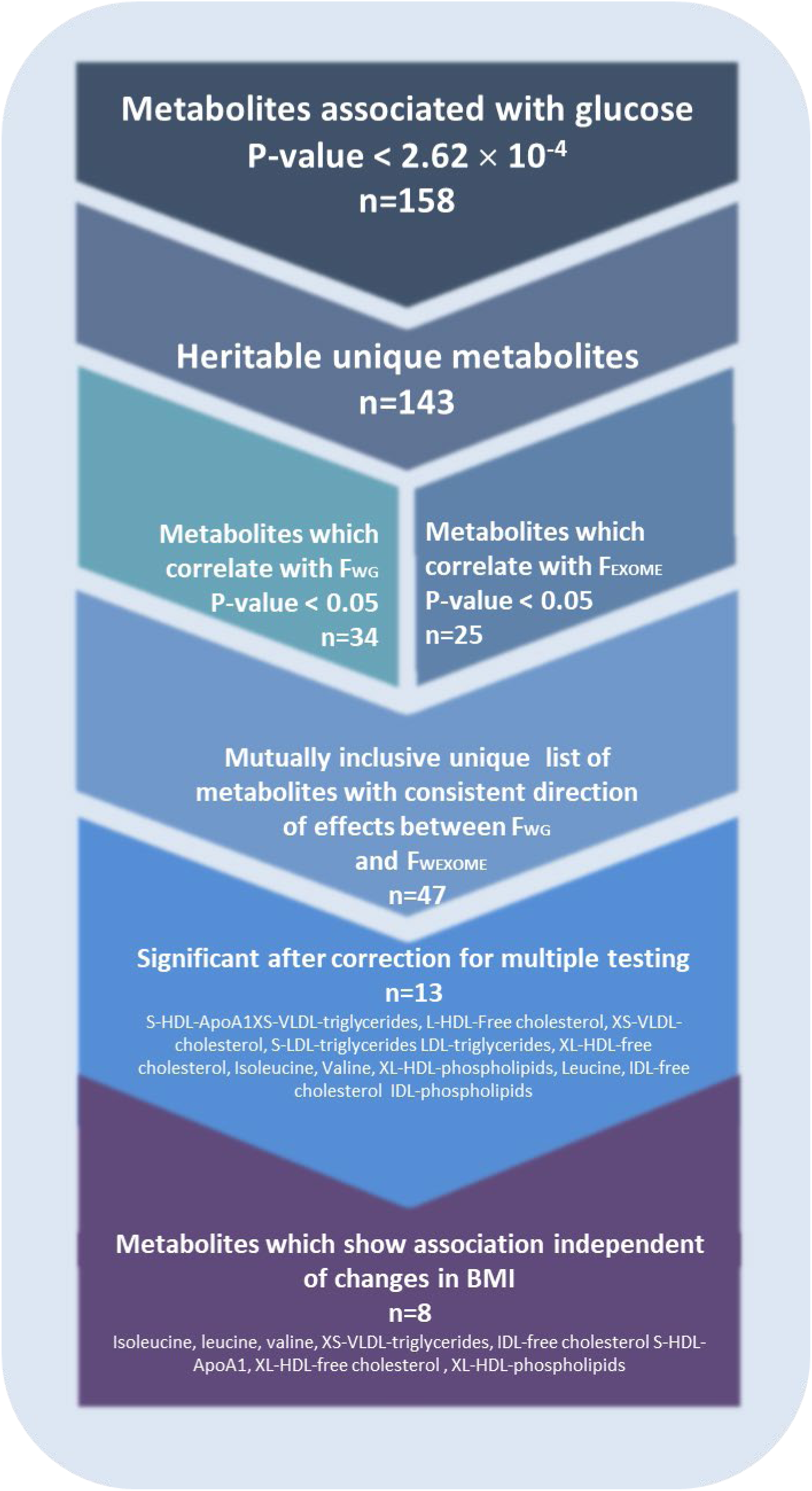
Study design.

## Methods

### Study population

The Erasmus Rucphen Family genetic isolate study (ERF) is a prospective family-based study located in Southwest of the Netherlands. This young genetic isolate was founded in the mid-eighteenth century and minimal immigration and marriages occurred between surrounding settlements due to social and religious reasons. The study population includes 3,465 individuals that are living descendants of 22 couples with at least six children baptized. Informed consent has been obtained from patients where appropriate. The study protocol was approved by the medical ethics board of the Erasmus Medical Center Rotterdam, the Netherlands^12^. The baseline demographic data and measurements of the ERF participants were collected around 2002 to 2006. All the participants filled out questionnaires on socio-demographics, diseases and medical history and lifestyle factors, and were invited to the research center for an interview and blood collection for biochemistry and physical examinations including blood pressure and anthropometric measurements have been performed. The participants were asked to bring all their current medications for registration during the interview. Venous blood samples were collected after at least eight hours of fasting. The detailed description of the ERF study and related measurements were reported previously^12^. Baseline type 2 diabetes was defined according to the fasting plasma glucose ≥ 7.0mmol/L and/or anti-diabetic treatment, yielding 212 cases and 2,564 controls, totaling up to 2,776 individuals. The follow-up data collection of the ERF study took place in May 2016 (9 to 14 years after baseline visit). During the follow-up, a total of 1,935 participants’ records were scanned for the incidence of type 2 diabetes in general practitioner’s databases. Additionally, a questionnaire on type 2 diabetes medication surveyed on 1,232 participants in June 2010 (4 to 8 years after baseline visit) was referred if a participant were not included in May 2016 follow-up. This effort yielded the inclusion of 18 otherwise missed extra cases, yielding a total of 349 cases and 2,427 controls included in the current study.

### Metabolite measurements and selection

#### Metabolite measurements

Metabolic markers were measured by five different metabolomics platforms using the methods which have been described in earlier publications^6,13–16^. In total 562 metabolic markers including sub-fractions of lipoproteins, triglycerides, phospholipids, ceramides, amino acids, acyl-carnitines and small intermediate compounds, which throughout this article will be referred as “*metabolites*”, were measured either by nuclear magnetic resonance (NMR) spectrometry or by mass spectrometry (MS). The platforms used in this research are: (1) Liquid Chromatography-MS (LC-MS, 116 positively charged lipids, comprising of 39 triglycerides, 47 phosphatidylcholines, 8 phosphatidylethanolamines, 20 sphingolipids, and 2 ceramides, available in up to 2,638 participants) measured in the Netherlands Metabolomics Center, Leiden using the method described before^13^; (2) Electrospray-Ionization MS (ESI-MS, in total 148 phospholipids and sphingolipids comprising of 16 plasmologens, 72 phosphatidylcholines, 27 phosphatidylethanolamines, 24 sphingolipids, 9 ceramides, available in up to 878 participants), measured in the Institute for Clinical Chemistry and Laboratory Medicine, University Hospital Regensburg, Germany using the method described previously^16^; (3) Small molecular compounds window based NMR spectroscopy (41 molecules comprising of 29 low-molecular weight molecules and 12 amino acids available in up to 2,639 participants) measured in the Center for Proteomics and Metabolomics, Leiden University Medical Center^6,17^; (4) Lipoprotein window-based NMR spectroscopy (104 lipoprotein particles size sub-fractions comprising of 28 VLDL components, 30 HDL components, 35 LDL components, 5 IDL components and 6 plasma totals, available in 2,609 participants) measured in the Center for Proteomics and Metabolomics, Leiden University Medical Center and lipoprotein sub-fraction concentrations were determined by the Bruker algorithm (Bruker BioSpin GmbH, Germany) as detailed in Kettunen *et al*^14^; (5) AbsoluteIDQTM p150 Kit of Biocrates Life Sciences AG (Biocrates, 153 molecules comprising of 14 amino acids, 91 phospholipids, 14 sphingolipids, 33 acyl-carnitines and hexose available in up to 989 participants) measured as detailed in publication from Draisma *et al*^*15*^ and the experiments were carried out at the Metabolomics Platform of the Genome Analysis Center at the Helmholtz Zentrum München, Germany as per the manufacturer’s instructions. The laboratories had no access to phenotype information.

### Genome-wide SNP measurements

Genotyping in ERF was performed using Illumina 318/370 K, Affymetrix 250 K, and Illumina 6K micro-arrays. All SNPs were imputed using MACH software (www.sph.umich.edu/csg/abecasis/MaCH/) based on the 1000G phase 1, release v3 reference. Individuals were excluded for excess autosomal heterozygosity, mismatches between called and phenotypic gender, and if there were outliers identified by an identical-by-state (IBS) clustering analysis.

### Exome sequence measurements

Exomes of 1,336 randomly selected individuals from the ERF study were sequenced at the Center for Biomics of the Cell Biology department, at the Erasmus Medical Center, the Netherlands. Sequencing was done at a median depth of 57× using the Agilent version V4 capture kit, on an Illumina Hiseq2000 sequencer, using the TruSeq Version 3 protocol. The sequence reads were aligned to the human genome build 19 (hg19), using Burrows-Wheeler Aligner (BWA) and the NARWHAL pipeline^18, 19 20^. Subsequently, the aligned reads were processed further, using the IndelRealigner, MarkDuplicates and Table Recalibration tools from the Genome Analysis Toolkit (GATK) ^21^ and Picard (http://picard.sourceforge.net). This was necessary to remove systematic biases and to recalibrate the PHRED quality scores in the alignments^22^. After processing, genetic variants were called, using the Unified Genotyper tool from the GATK^21^. For each sample, at least 4 Gigabases of sequence was aligned to the genome. Functional annotations were also performed using the SeattleSeq annotation 138 database. About 1.4 million SNVs were called. After removing variants with low quality, out of Hardy-Weinberg equilibrium (HWE, P-value < 10^−6^) and low call rate (< 99%), and samples with a low call rate (< 90%), we retrieved 543,954 very high-quality SNVs in 1,327 individuals.

### Exome chip measurements

Study participants from the ERF study whose exomes were not sequenced (N = 1,527) were genotyped on the Illumina Infinium HumanExome BeadChip, version 1.1, which contains over 240,000 exonic variants selected from multiple sources together spanning 12,000 samples from multiple ethnicities. Calling was performed with GenomeStudio. We removed subjects with a call rate < 95%, IBS > 0.99 and heterozygote ratio > 0.60. Ethnic outliers identified using a principal component analysis with 1000 Genomes data and individuals with sex discrepancies were also removed. The SNVs that were monomorphic in our sample or had a call rate < 95% were removed. After quality control, we retrieved about 70,000 polymorphic SNVs in 1,515 subjects.

### Statistical methods

The outlying metabolite values that were more than four times standard deviation away from the mean were excluded from the analysis. Non-normally distributed measurements were natural logarithm transformed, or rank transformed accordingly. As described previously^23^, in brief, we assessed the pairwise partial correlation between each metabolite and each glycemic trait (i.e., fasting glucose, fasting insulin, homeostatic model assessment for insulin resistance (HOMA-IR), BMI and WHR) in the non-diabetic participants. The association of metabolites and T2DM were assessed by logistic regression with T2DM status as the dependent variable. We included age, sex and lipid-lowering medication as covariates and adjusted the models by familial relatedness by using the polygenic residuals extracted by the “polygenic” function of R package GenABEL. We applied a Bonferroni correction based on the number of independent vectors in the data which were estimated to be 191 by Matrix Spectral Decomposition (MSD)from the 562 metabolites^24^. A P-value < 2.62 × 10^−4^ (0.05/191) was used as the threshold for metabolome-wide significance, yielding 158 metabolites associated with glucose. Among those, 124 of them have already been shown by us in an earlier report^23^. **Supplementary Table 1** shows the metabolites and their association with glucose, BMI, WHR, insulin, HOMA-IR and T2DM. Seven metabolites (Phosphatidylcholine alkyl-acyl 34:3, Phosphatidylcholine alkyl-acyl 36:2, Phosphatidylcholine alkyl-acyl 44:5, Phosphatidylcholine diacyl 38:4, Phosphatidylcholine diacyl 40:4, Valine and Tyrosine) were measured independently by more than one platforms (i.e. LC-MS, ESI-MS, Biocrates) and we discarded the measurement with smaller sample size. As a result, 151 metabolites were taken forward to the next step.

We estimated the heritability (H^2^) for the 151 unique metabolites as well as fasting glucose as a reference in ERF pedigree using SOLAR software^25^ with age and sex adjusted. 143 of the tested metabolites showed a significant heritable component (P-value < 0.05) and were taken to the next step. We calculated the inbreeding coefficient (F) using whole genome array SNPs (F_*WG*_), SNVs from exome sequence (F_*WXSEQ*_) and exome array (F_*WXCHIP*_) per individual using the --het function in PLINK software^26^. Pedigree-based inbreeding coefficient (F_*PED*_) was also calculated based on identical by descent (IBD) sharing. Five individuals, four measured by exome array and one by exome sequence, were excluded due to them lying outside of the normal distribution of the F values. The genotype data of a population-based cohort, Rotterdam Study^27^ (n = 6,291), was also used when comparing the inbreeding coefficient between family-based study and population study. We calculated the Pearson’s correlation coefficients between inbreeding and each metabolite (143 metabolites) and risk factors of T2DM adjusted first by familial relatedness, age, sex and lipid-lowering medication and in a second model, we additionally adjusted for BMI. The summary statistics deriving from exomic coefficients (F_*WXSEQ*_ and F_*WXCHIP*_) coming from independent datasets were meta-analyzed in order to increase the sample size (F_*EXOME*_) using the R package Metafor, after excluding the overlapping 18 individuals at random from either of the sets. For correlation analysis of the pedigree-based F_*PED*_ which was not normally distributed, Spearman’s R was used. Genome-wide association study (GWAS) of the candidate metabolites were performed by using *--all models* and --*mmscore* functions as implemented in the ProbABEL software (v.0.3.0), using 1000G imputation based genetic data of the ERF population, adjusted by age and sex, additionally correcting for relatedness using mixed model approach.

## Results

### Heritability estimations

**Figure 2** shows the H^2^ estimates and the effects of the shared environment for the 143 significantly heritable metabolites as clustered in chemical groups of small molecular compounds and amino acids, phospholipids and sphingolipids, triglycerides and lipoproteins, as well as fasting glucose as a reference. The H^2^ was estimated to be highest for lipoprotein group, remarkably for L-HDL-Free cholesterol (H^2^ =0.39, P-value =6.9 × 10^−30^), HDL-Free cholesterol (H^2^ =0.37, P-value =9.6 × 10^−28^), L-HDL-cholesterol (H^2^ =0.35, P-value =3.9 × 10^−23^) and XL-HDL-phospholipids (H^2^ =0.34, P-value =3.0 × 10^−17^). The H^2^ was estimated to be lowest for some poly-unsaturated phospholipids such as: Phosphatidylcholine diacyl 40:6 (H^2^ =0.058, P-value =0.017), Phosphatidylcholine diacyl 34:4 (H^2^ =0.063, P-value =0.016), Phosphatidylcholine acyl-alkyl 44:6 (H^2^ =0.065, P-value =0.007) and Triglycerides (46:2) (H^2^ =0.059, P-value =0.026). The full list is given in **Supplementary Table 2**. The 143 metabolites significantly heritable were taken to the next step.

**Figure 2.**
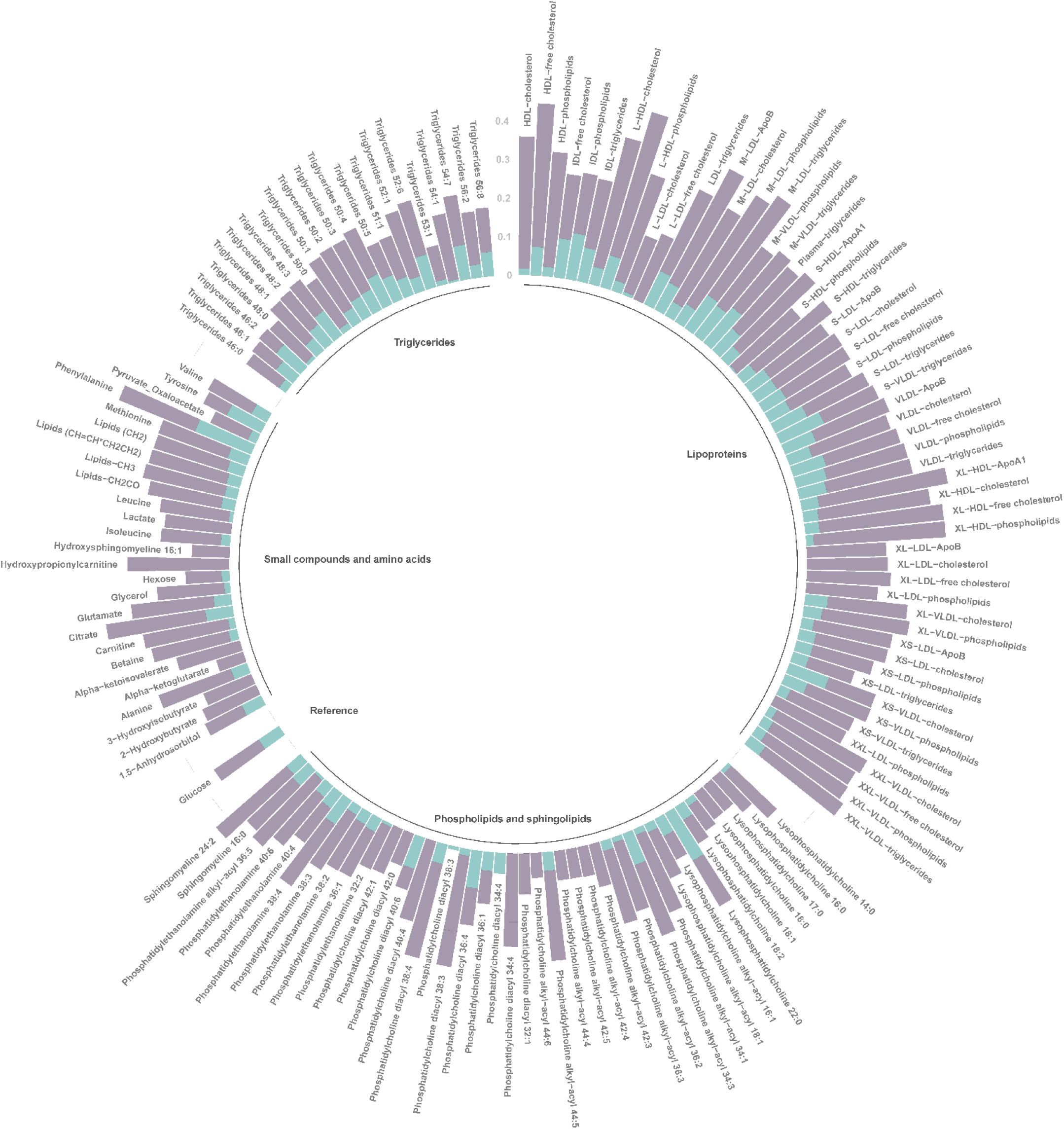
Influence of genetic factors and shared environmental factors on the traits studied. **Purple**: genetic heritability (H^2^). **Green**: effects of the shared environment.

### Correlation of inbreeding with T2DM metabolites

The inbreeding coefficients for ERF population and their comparisons to the ones calculated for the (outbred) Rotterdam Study population are given in **Figure 3A**. The family-based ERF study has much higher skewness value than the population-based Rotterdam study. The separation between the distributions is the most remarkable for the F_*EXOME*_ values with skewness = 0.85 in ERF study and skewness = 0.16 in the Rotterdam Study. **Figure 3B** shows the correlation between the F values based on the whole genome, exome or pedigree (n = 2,916). It shows high correlation among the three inbreeding coefficient methods with correlation coefficient ranging from 0.37 (correlation of F_*PED*_ and F_*EXOME*_, P-value = 1.8 × 10^−76^) to 0.59 (correlation for F_*PED*_ and F_*WG*_, P-value = 2.0 × 10^−271^). **Figure 4** shows the heatmap generated based on the significant correlation between the inbreeding coefficients and metabolites and glycemic traits (i.e., T2DM, fasting glucose, fasting insulin, HOMA-IR, BMI and WHR). We found the inbreeding coefficients derived by either method correlated significantly and positively with BMI, WHR, glucose, insulin and HOMA-IR as a measure of insulin resistance (P-value < 0.05, **Supplementary Table 3** and **Figure 4**). The association of BMI with F_*WG*_ is the strongest one (r = 0.09, P-value = 8.5 × 10^−6^). Among the selected 143 metabolites, in total 47 metabolites correlated with at least one of the inbreeding coefficients when adjusting for age, sex, and lipid-lowering medication: 34 correlated with F_*WG*_, 25 with F_*EXOME*_, and one with F_*PED*_ with a suggestive P-value < 0.05 (**Supplementary Table 4**).

**Figure 3A.**
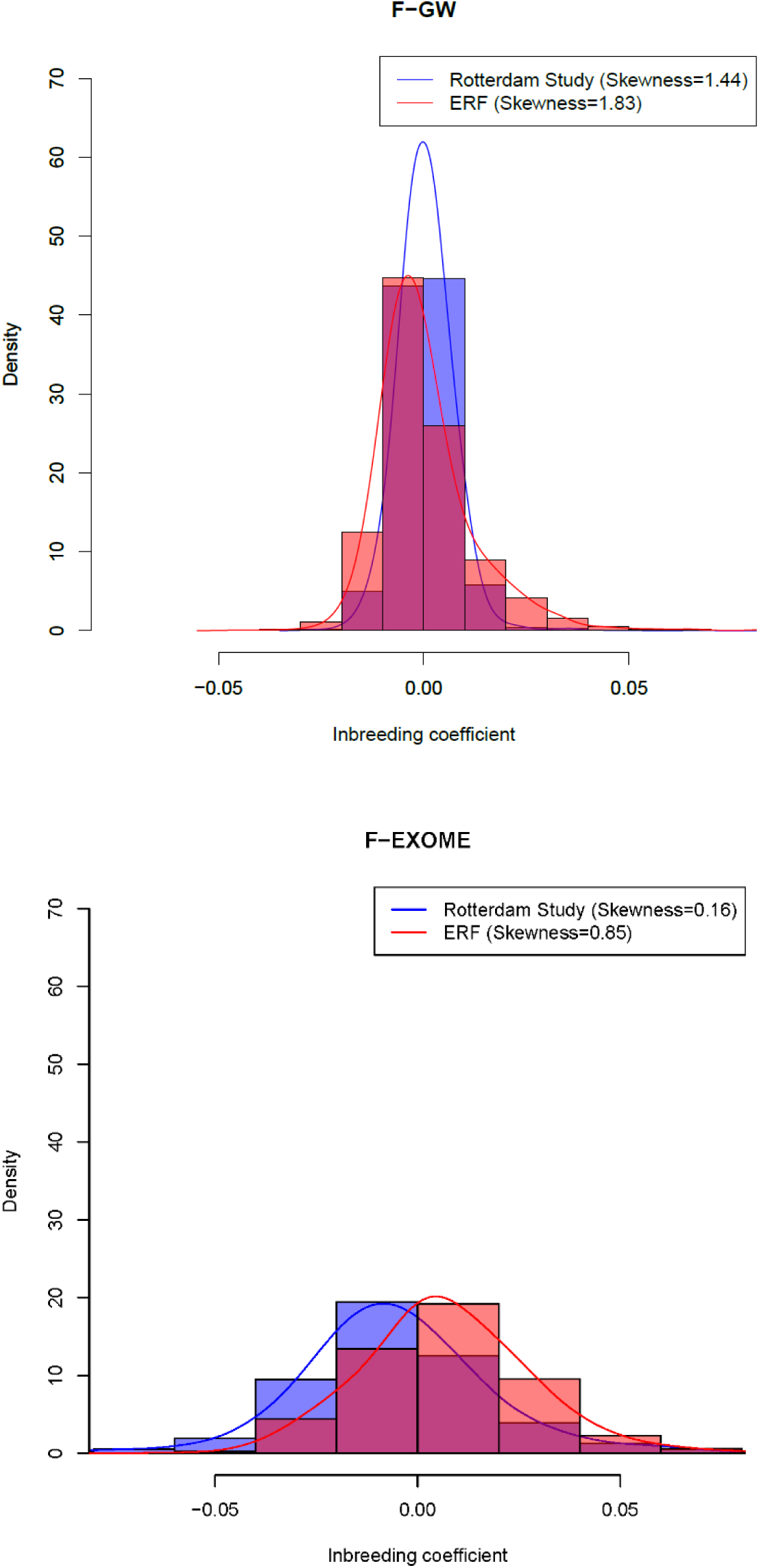
Comparison histogram plots of inbreeding coefficients of the consanguineous and the example outbred population. F-GW: whole genome array based inbreeding coefficient. F-EXOME: the combination of exome chip and exome sequence based inbreeding coefficient.

**Figure 3B.**
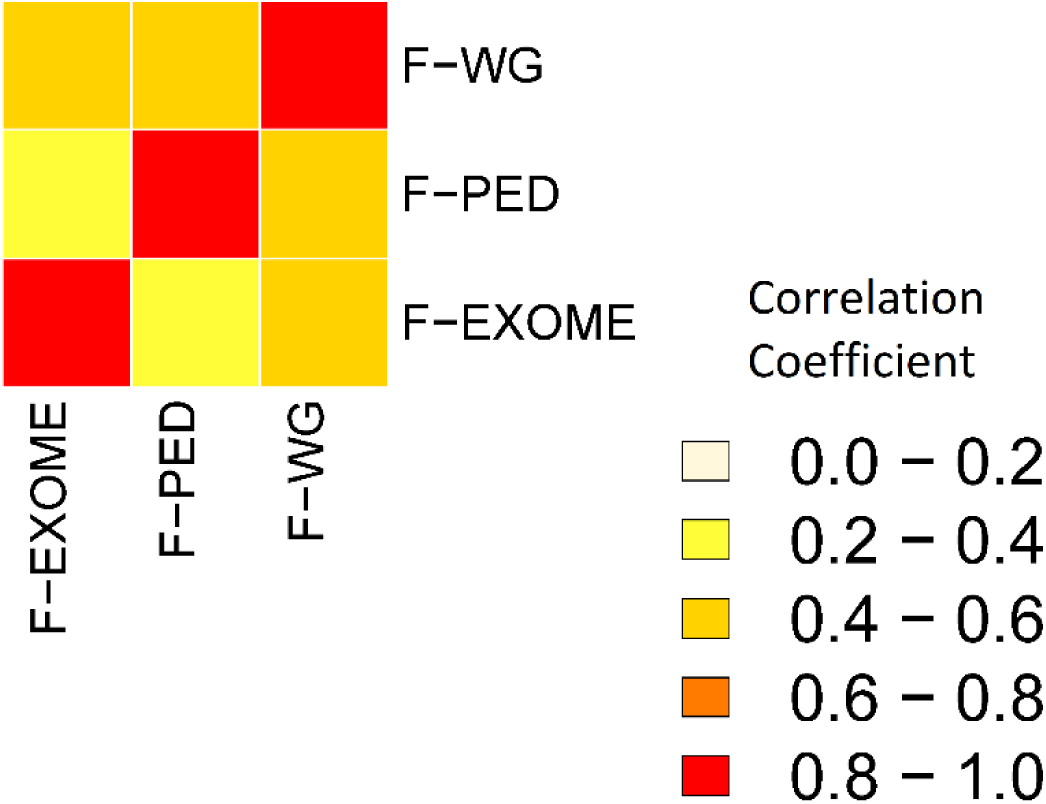
Similarity based on correlation coefficient between the inbreeding values calculated within the ERF population. F-GW: whole genome array based inbreeding coefficient. F-EXOME: the combination of exome chip and exome sequence based inbreeding coefficient. F-PED: Pedigree based inbreeding coefficient.

**Figure 4.**
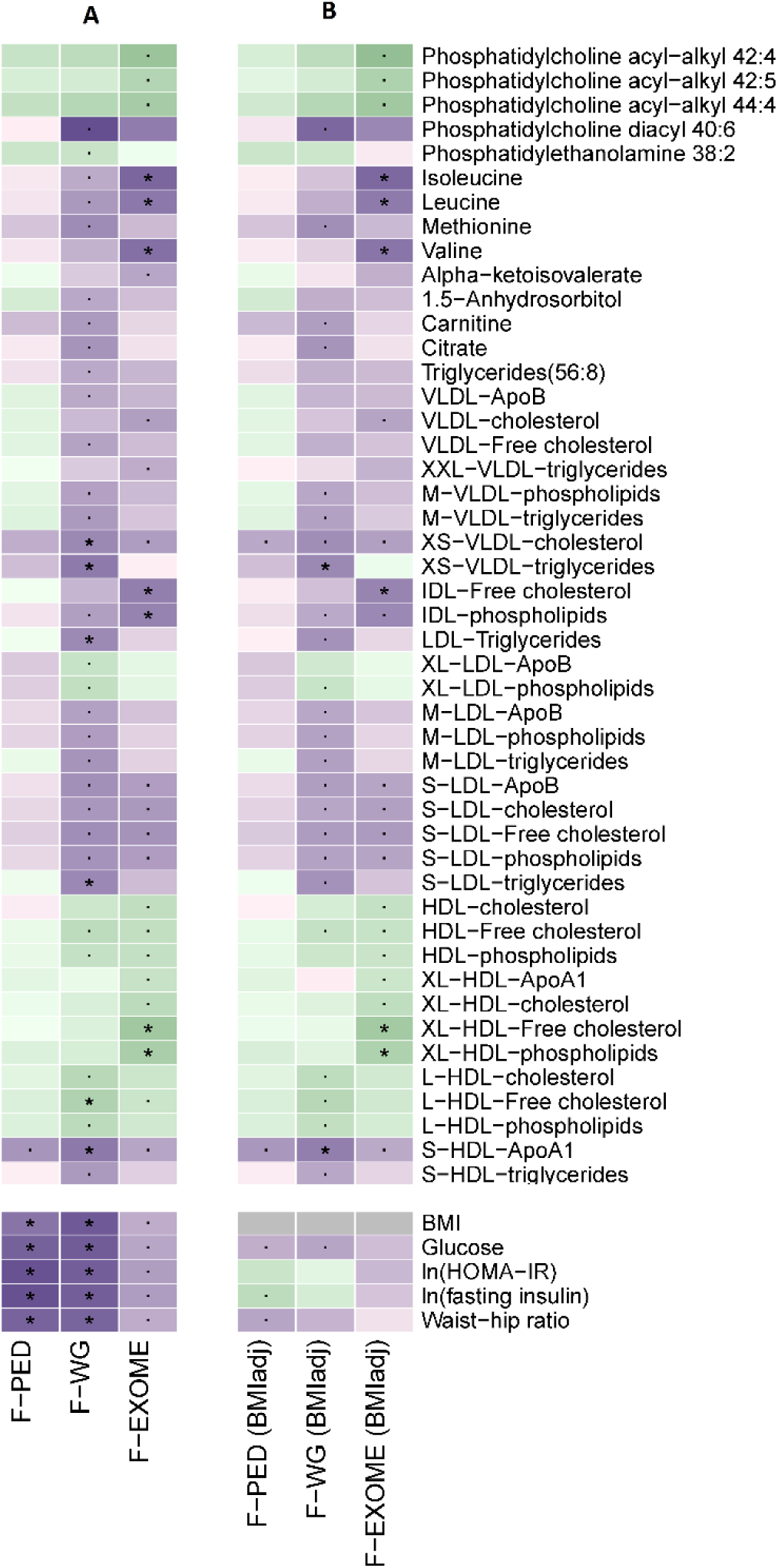
The effect of inbreeding values on the metabolite profiles and relation to T2DM risk factors. **Figure 4A** shows the correlation between the inbreeding coefficients and the metabolites and T2DM-related risk factors. **Figure 4B** correlation between the inbreeding coefficients and the metabolites after adjusting for BMI. Metabolites with at least one suggestive P-value of association (<0.05) are shown with “.”. Significance after correction for multiple testing (P-value < 2.8 × 10^−3^) is marked with a “* “. **Purple**: positive association. **Green**: negative association. The depth of purple and green presents the value of correlation coefficient. F-GW: whole genome array based inbreeding coefficient. F-EXOME: the combination of exome chip and exome sequence based inbreeding coefficient. F-PED: Pedigree based inbreeding coefficient.

Among those 12 metabolites were common (P-value < 0.05) in both whole genome-based and exome-based analyses. These were: BCAAs; leucine and isoleucine, HDL fractions; L-HDL-free cholesterol, S-HDL-ApoA1, HDL-free cholesterol, HDL-phospholipids, LDL fractions; S-LDL-ApoB, S-LDL-cholesterol, S-LDL-free cholesterol, S-LDL-phospholipids, and larger particle fractions; IDL-phospholipids and XS-VLDL-cholesterol. When corrected for the multiple testing (P-value < 2.8 × 10^−3^ based on the 18 independent vectors tested in the 47 metabolites), six metabolites remained significantly correlated with F_*WG*_ (S-HDL-ApoA1, XS-VLDL-triglycerides, L-HDL-Free cholesterol, XS-VLDL-cholesterol, S-LDL-triglycerides and LDL-triglycerides) and seven with F_*EXOME*_ (XL-HDL-free cholesterol, Isoleucine, Valine, XL-HDL-phospholipids, Leucine, IDL-free cholesterol and IDL-phospholipids). The strongest association was observed with S-HDL-ApoA1 (r= 0.07, P-value= 4.5 × 10^−4^ for F_*WG*)_. S-HDL-ApoA1 also correlated with F_*PED*_ (r= 0.05, P-value= 8.0 × 10^−3^) and F_*EXOME*_ (r= 0.04, P-value= 0.024) (**Supplementary Table 4**). As BMI is well known to be influenced by both homozygosity of the individuals and T2DM, we next tested if the observed associations are due to the differences in BMI. Thirty-nine of the 47 metabolites correlated with at least one of the inbreeding coefficients when adjusting for BMI in addition (**Supplementary Table 5**). Eight metabolites remained statistically significant after correcting for multiple testing (P-value < 2.8 × 10^−3^), including S-HDL-ApoA1 and XS-VLDL-triglycerides with F_*WG*_ and IDL-free cholesterol, Isoleucine, Leucine, Valine, XL-HDL-free cholesterol and XL-HDL-phospholipids with F_*EXOME*_ when we accounted for BMI. Six of the metabolites were positively correlated with inbreeding coefficients. These were isoleucine (r= 0.08, P-value= 3.1 × 10^−4^), leucine (r= 0.07, P-value= 1.6 × 10^−3^), valine (r= 0.07, P-value= 9.7 × 10^−4^), XS-VLDL-triglycerides (r= 0.06, P-value= 2.2 × 10^−3^), IDL-free cholesterol (r= 0.06, P-value= 2.3 × 10^−3^) and S-HDL-ApoA1 (r= 0.07, P-value= 9.7 × 10^−4^). Two extra-large HDL related measurements correlated negatively with the inbreeding coefficient. These were XL-HDL-free cholesterol (r= −0.08, P-value= 2.7 × 10^−4^) and XL-HDL-phospholipids (r= −0.07, P-value= 1.4 × 10^−3^). Overall the correlations showed a consistent pattern across the three different inbreeding coefficients. Full results from the BMI adjusted model are given in **Supplementary Table 5.**

### Model supervised analysis of known genetic determinants

As we found 13 unique metabolites significantly correlated with either inbreeding coefficient after adjustment of age, sex and lipid-lowering medication, we directly checked if the known genetic determinants of these 13 metabolites could yield to smaller type 1 error for rejecting the null hypothesis by genetic models taking the recessive inheritance into account. Nine out of the 13 metabolites (Isoleucine, Leucine, Valine, IDL-free cholesterol, IDL-phospholipids, L-HDL-free cholesterol, XL-HDL-free cholesterol, XL-HDL-phospholipids and XS-VLDL-triglycerides) were studied by GWAS before^14^. For those we selected, in total 6,379 SNP-metabolite pairs which have been previously reported with P-value < 5 × 10^−8^ using the additive genetic model (n = 24,925). Out of the list of 6,379 pairs, we identified 189 occurrences in which resulted in at least 10 times reduction in type 1 error (one time the P-value gain in log scale) as compared to the additive model analyzing the same SNP-metabolite pair in ERF data (**Supplementary Table 6**). These associations were detected for SNPs in genomic regions harboring known metabolism genes *PPM1K, LIPC, USP24, LPL, APOB, ABCG5, PLTP* and *APOC1* (the full list is given in **Supplementary Table 6**).

The remaining four metabolites (LDL-triglycerides, S-HDL-ApoA1, S-LDL-triglycerides and XS-VLDL-cholesterol) were not studied in GWAS before and for those we performed GWAS in ERF. For S-HDL-ApoA1, we successfully detected a novel region located in 1q41 by using the genetic model that tests the dominant effect of the effect allele used (hence, the recessive model for the non-effect allele). In total three SNPs (rs78230510, rs118043948 and rs1538560) in the vicinity of *DISP1* gene passed the predefined genome-wide significance threshold for the dominant model (P-value < 5 × 10^−8^, **Supplementary Table 7**). The GWAS of the remaining traits did not yield any novel genome-wide significant loci, neither performed better in additive, recessive or over-dominant genetic models.

## Discussion

Focusing on 143 T2DM related metabolites in a genetically isolated population, we detected a strong heritable component and evidence for the effects of inbreeding on metabolism at the population level. We showed that some part of the variation in all of the metabolites studied can be attributed to the pedigree component, with heritability ranging from 0.06 to 0.39. We also showed that a total of 13 metabolites are influenced by inbreeding, among which S-HDL-ApoA1 (ApoA1 content of the small HDL lipoprotein) shows a robust positive correlation with inbreeding coefficient estimated based on all of the three independent genetic measurements. We also showed that for isoleucine, leucine, valine, XS-VLDL-triglycerides, IDL-free cholesterol, S-HDL-ApoA1, XL-HDL-free cholesterol and XL-HDL-phospholipids, the effect of inbreeding was independent of BMI. These results show that T2DM related metabolic profile based on lipoproteins and amino-acids that is present in clinically healthy individuals is influenced by the excess of homozygosity of the individuals.

Consanguinity, i.e., mating between related individuals, results in increased probability to inherit two identical copies of the same ancestral gene in the offspring and thus an increased risk of recessive disease. Harmful effects of close consanguinity in humans have been shown for several of outcomes including intelligence^28^, schizophrenia^29^, bipolar disorder^30^, hypertension^31^, heart disease^32^, cancer^33^, but notably also for metabolic health^34^ and T2DM^4^. Since consanguinity is strongly associated with risk for multifactorial disease, it is more than likely that unknown homozygous regions of the genome explain a significant portion of familial aggregation among other possible mechanisms^35^. Runs of homozygosity (ROHs) have been associated with several human traits such as personality^36^, schizophrenia^37^, short stature^5^ and birth height^3^. To our knowledge, this has been the first report showing the impact of genome-wide recessive effects on metabolomics at the population level. Our current results from in-depth metabolomics analysis of plasma lipids are complementary to the earlier report which has already shown the increasing effect of inbreeding on commonly measured plasma lipid levels of triglycerides, total cholesterol and low-density lipoprotein cholesterol (LDL-C)^34^ in the ERF population. Using a high-resolution metabolic panel, we additionally show that S-HDL-ApoA1, XS-VLDL-triglycerides, IDL-free cholesterol, XL-HDL-free cholesterol and XL-HDL-phospholipids are also under control of inbreeding. Overall, inbreeding has a positive effect on unfavorable metabolic measures, for HDL lipidome a more complex scenario exists, depending on particle size. Previously we have shown the heterogenous signals of the HDL lipidome in relation to metabolic health; the decreasing effect of HDL-cholesterol on fasting glucose is more specific to the large and extra-large particle subclasses, whereas for small HDL component, an increasing effect exists^23^. The genetic determinants have been identified for these metabolites with variable success depending on the genetic architecture.

A recent GWAS identified SNPs that explain variance 0.5% for isoleucine, 0.7% for leucine, 10% for valine, 7% for XS-VLDL-triglycerides, 11% in IDL-free cholesterol, 7% in XL-HDL-free cholesterol and 9 % in XL-HDL-phospholipids. Whereas S-HDL-ApoA1 was not covered by large GWAS yet^14^. The fact that the GWAS explained only a small part of the variation using an additive genetic model, could be explained by that there is a substantial recessive component of the genetic architecture to be investigated. We further tested these top genetic determinants in non-additive models and found that indeed within the same loci there is evidence for the association which yields higher power of detection in recessive or dominants models rather than the additive model. However, it was only the case for a limited proportion of top SNPs analyzed, located in the vicinity of metabolic genes *PPM1K, LIPC, USP24, LPL, APOB, ABCG5, PLTP* and *APOC1*. We additionally performed a GWAS in much smaller sample size on S-HDL-ApoA1 and we identified three genetic variants within the vicinity of *DISP1* gene. The top SNP rs78230510 with minor allele frequency (MAF) of 6% is located within a promoter flanking region and the second top SNP rs118043948 is located intronic within the *DISP1* gene. The third SNP has MAF of 17% in ERF and is also located in the intron of *DISP1*. A phenome-wide association study (PheWAS) look-up for these SNPs in GWASATLAS by Watanabe K et al^38^, including 4141 additive model based GWAS tables, did not yield any significant associations at the genome-wide level.

One limitation of our study is that S-HDL-ApoA1 is not measured in any other cohort to our knowledge, giving no chances of replication. Model supervised GWAS for metabolic measurements has been very limited by one report only from Tsepilov YA et al^39^, who have GWAS of metabolites from Biocrates platform only, covering mainly the phospholipids pathways. To our knowledge remaining promising metabolic pathways are yet to be studied.

In conclusion, our study points out several metabolic pathways, clustered under the metabolic groups of lipoproteins, amino-acids and phosphatidylcholines related to T2DM are partially under recessive genetic control. The genetic association analysis of these metabolites in the recessive model is suggested to be performed in larger sample size studies.

## Acknowledgments

ERF was supported by the Consortium for Systems Biology (NCSB), both within the framework of the Netherlands Genomics Initiative (NGI)/Netherlands Organization for Scientific Research (NWO). ERF study as a part of EUROSPAN (European Special Populations Research Network) was supported by European Commission FP6 STRP grant number 018947 (LSHG-CT-2006-01947) and also received funding from the European Community’s Seventh Framework Program (FP7/2007-2013)/grant agreement HEALTH-F4-2007-201413 by the European Commission under the program “Quality of Life and Management of the Living Resources” of 5th Framework Program (no. QLG2-CT-2002-01254) as well as FP7 project EUROHEADPAIN (nr 602633). High-throughput analysis of the ERF data was supported by joint grant from Netherlands Organization for Scientific Research and the Russian Foundation for Basic Research (NWO-RFBR 047.017.043). High throughput metabolomics measurements of the ERF study has been supported by BBMRI-NL (Biobanking and Biomolecular Resources Research Infrastructure Netherlands). Ayse Demirkan is supported by a Veni grant (2015) from ZonMw. Ayse Demirkan, Jun Liu and Cornelia van Duijn have used exchange grants from the Personalized pREvention of Chronic DIseases consortium (PRECeDI) (H2020-MSCA-RISE-2014). The funders had no role in study design, data collection and analysis, decision to publish, or preparation of the manuscripts. ERF study is grateful to all study participants and their relatives, general practitioners and neurologists for their contributions and to P. Veraart for her help in genealogy, J. Vergeer for the supervision of the laboratory work and P. Snijders for his help in data collection.

## Author Contributions

A.D, J.L. and C.M.v.D contributed to study design. K.W.v.D. and C.M.v.D contributed to data collection. A.D., J.L. and N.A. contributed to data analysis. A.D., J.L. contributed to writing of manuscript. All the authors contributed to critical review of manuscript.

## References

1 Vernon, H. J. Inborn Errors of Metabolism: Advances in Diagnosis and Therapy. JAMA Pediatr 169, 778–782, doi:2323438 [pii] 10.1001/jamapediatrics.2015.0754 (2015).

2 Howrigan, D. P. et al. Genome-wide autozygosity is associated with lower general cognitive ability. Mol Psychiatry 21, 837–843, doi:mp2015120 [pii] 10.1038/mp.2015.120 (2016).

3 Verweij, K. J. et al. The association of genotype-based inbreeding coefficient with a range of physical and psychological human traits. PLoS One 9, e103102, doi:10.1371/journal.pone.0103102 PONE-D-14-19464 [pii] (2014).

4 Gosadi, I. M., Goyder, E. C. & Teare, M. D. Investigating the potential effect of consanguinity on type 2 diabetes susceptibility in a saudi population. Hum Hered 77, 197–206, doi:000362447 [pii] 10.1159/000362447 (2014).

5 McQuillan, R. et al. Evidence of inbreeding depression on human height. PLoS Genet 8, e1002655, doi:10.1371/journal.pgen.1002655 PGENETICS-D-12-00175 [pii] (2012).

6 Demirkan, A. et al. Insight in genome-wide association of metabolite quantitative traits by exome sequence analyses. PLoS Genet 11, e1004835, doi:10.1371/journal.pgen.1004835 (2015).

7 Merino, J. et al. Metabolomics insights into early type 2 diabetes pathogenesis and detection in individuals with normal fasting glucose. Diabetologia 61, 1315–1324, doi:10.1007/s00125-018-4599-x (2018).

8 Kahn, S. E., Cooper, M. E. & Del Prato, S. Pathophysiology and treatment of type 2 diabetes: perspectives on the past, present, and future. Lancet 383, 1068–1083, doi:10.1016/S0140-6736(13)62154-6 (2014).

9 Wang, C. et al. Plasma phospholipid metabolic profiling and biomarkers of type 2 diabetes mellitus based on high-performance liquid chromatography/electrospray mass spectrometry and multivariate statistical analysis. Anal Chem 77, 4108–4116, doi:10.1021/ac0481001 (2005).

10 Wang, T. J. et al. Metabolite profiles and the risk of developing diabetes. Nat Med 17, 448–453, doi:10.1038/nm.2307 (2011).

11 Liu, J. et al. Metabolomics based markers predict type 2 diabetes in a 14-year follow-up study. Metabolomics 13, 104, doi:10.1007/s11306-017-1239-2 1239 [pii] (2017).

12 Santos, R. L. et al. Heritability of fasting glucose levels in a young genetically isolated population. Diabetologia 49, 667–672, doi:10.1007/s00125-006-0142-6 (2006).

13 Gonzalez-Covarrubias, V. et al. Lipidomics of familial longevity. Aging Cell 12, 426–434, doi:10.1111/acel.12064 (2013).

14 Kettunen, J. et al. Genome-wide study for circulating metabolites identifies 62 loci and reveals novel systemic effects of LPA. Nat Commun 7, 11122, doi:ncomms11122 [pii] 10.1038/ncomms11122 (2016).

15 Draisma, H. H. et al. Genome-wide association study identifies novel genetic variants contributing to variation in blood metabolite levels. Nat Commun 6, 7208, doi:10.1038/ncomms8208 (2015).

16 Demirkan, A. et al. Genome-wide association study identifies novel loci associated with circulating phospho- and sphingolipid concentrations. PLoS Genet 8, e1002490, doi:10.1371/journal.pgen.1002490 (2012).

17 Verhoeven, A., Slagboom, E., Wuhrer, M., Giera, M. & Mayboroda, O. A. Automated quantification of metabolites in blood-derived samples by NMR. Analytica Chimica Acta (2017).

18 Li, H. & Durbin, R. Fast and accurate short read alignment with Burrows-Wheeler transform. Bioinformatics 25, 1754–1760, doi:btp324 [pii] 10.1093/bioinformatics/btp324 (2009).

19 Brouwer, R. W., van den Hout, M. C., Grosveld, F. G. & van Ijcken, W. F. NARWHAL, a primary analysis pipeline for NGS data. Bioinformatics 28, 284–285, doi:btr613 [pii] 10.1093/bioinformatics/btr613 (2012).

20 Li, H. et al. The Sequence Alignment/Map format and SAMtools. Bioinformatics 25, 2078–2079, doi:btp352 [pii] 10.1093/bioinformatics/btp352 (2009).

21 McKenna, A. et al. The Genome Analysis Toolkit: a MapReduce framework for analyzing next-generation DNA sequencing data. Genome Res 20, 1297–1303, doi:gr.107524.110 [pii] 10.1101/gr.107524.110 (2010).

22 Ewing, B. & Green, P. Base-calling of automated sequencer traces using phred. II. Error probabilities. Genome Res 8, 186–194 (1998).

23 Liu, J. et al. A Mendelian Randomization Study of Metabolite Profiles, Fasting Glucose, and Type 2 Diabetes. Diabetes 66, 2915–2926, doi:db17-0199 [pii] 10.2337/db17-0199 (2017).

24 Li, J. & Ji, L. Adjusting multiple testing in multilocus analyses using the eigenvalues of a correlation matrix. Heredity (Edinb) 95, 221–227, doi:6800717 [pii] 10.1038/sj.hdy.6800717 (2005).

25 Almasy, L. & Blangero, J. Multipoint quantitative-trait linkage analysis in general pedigrees. Am J Hum Genet 62, 1198–1211, doi:S0002-9297(07)61542-0 [pii] 10.1086/301844 (1998).

26 Purcell, S. et al. PLINK: a tool set for whole-genome association and population-based linkage analyses. Am J Hum Genet 81, 559–575, doi:10.1086/519795 (2007).

27 Ikram, M. A. et al. The Rotterdam Study: 2018 update on objectives, design and main results. Eur J Epidemiol 32, 807–850, doi:10.1007/s10654-017-0321-4 10.1007/s10654-017-0321-4 [pii] (2017).

28 Fareed, M. & Afzal, M. Estimating the inbreeding depression on cognitive behavior: a population based study of child cohort. PLoS One 9, e109585, doi:10.1371/journal.pone.0109585 PONE-D-14-23257 [pii] (2014).

29 Mansour, H. et al. Consanguinity and increased risk for schizophrenia in Egypt. Schizophr Res 120, 108–112, doi:S0920-9964(10)01197-7 [pii] 10.1016/j.schres.2010.03.026 (2010).

30 Mansour, H. et al. Consanguinity associated with increased risk for bipolar I disorder in Egypt. Am J Med Genet B Neuropsychiatr Genet 150B, 879–885, doi:10.1002/ajmg.b.30913 (2009).

31 Rudan, I. et al. Inbreeding and the genetic complexity of human hypertension. Genetics 163, 1011–1021 (2003).

32 Shami, S. A., Qaisar, R. & Bittles, A. H. Consanguinity and adult morbidity in Pakistan. Lancet 338, 954, doi:0140-6736(91)91828-I [pii] (1991).

33 Lebel, R. R. & Gallagher, W. B. Wisconsin consanguinity studies. II: Familial adenocarcinomatosis. Am J Med Genet 33, 1–6, doi:10.1002/ajmg.1320330102 (1989).

34 Isaacs, A. et al. Heritabilities, apolipoprotein E, and effects of inbreeding on plasma lipids in a genetically isolated population: the Erasmus Rucphen Family Study. Eur J Epidemiol 22, 99–105, doi:10.1007/s10654-006-9103-0 (2007).

35 Hoppenbrouwers, I. A. et al. Familial clustering of multiple sclerosis in a Dutch genetic isolate. Mult Scler 13, 17–24 (2007).

36 Verweij, K. J. et al. Maintenance of genetic variation in human personality: testing evolutionary models by estimating heritability due to common causal variants and investigating the effect of distant inbreeding. Evolution 66, 3238–3251, doi:10.1111/j.1558-5646.2012.01679.x (2012).

37 Keller, M. C. et al. Runs of homozygosity implicate autozygosity as a schizophrenia risk factor. PLoS Genet 8, e1002656, doi:10.1371/journal.pgen.1002656 PGENETICS-D-11-02271 [pii] (2012).

38 Watanabe, K. et al. A global view of pleiotropy and genetic architecture in complex traits. bioRxiv, 500090 (2018).

39 Tsepilov, Y. A. et al. Nonadditive Effects of Genes in Human Metabolomics. Genetics 200, 707–718, doi:10.1534/genetics.115.175760 (2015).

